# A fluorescent reporter and single-turnover kinetics reveal new insight into BAM complex function

**DOI:** 10.1101/2025.05.30.657120

**Authors:** Whitney N. Bergman, Marcelo C. Sousa

## Abstract

The β-barrel assembly machine (BAM) is an essential protein complex that folds and inserts outer membrane β-barrel proteins (OMPs) into the bacterial outer membrane. The BAM complex contains the essential BamA OMP with five soluble polypeptide transport-associated (POTRA) domains, which scaffold the essential BamD lipoprotein and the non-essential lipoproteins BamB/C/E. The importance of each BAM component has been investigated primarily using cell-based phenotypic assays, and structural data have revealed insights into the role of the BamA β-barrel in OMP folding. However, *in vitro* quantitative analysis for the function of each BAM component has been challenging. We describe the development of a fluorescent reporter of OMP folding, tOmpA-A488, which we use to obtain single-turnover kinetic parameters for wildtype BAM complex *in vitro*. We observe a k_fold_ of 0.78 ± 0.15 min^−1^ and approximate substrate affinity of 3.1 ± 1.1 µM consistent with estimates of *in vivo* requirements. We also find that, contrary to prevailing models, POTRA domain deletions that include POTRA3, which is essential in cells, do not drastically impact activity. This indicates that the first three POTRA domains do not play a major role in binding or folding OMPs under single-turnover conditions, suggesting a different role in cells. Furthermore, we find that BamA alone is inactive in *E. coli* lipid liposomes, and the gain-of-function mutant BamA E470K does not rescue activity *in vitro*. The single-turnover kinetics enabled by the fluorescent reporter presented here defines a robust platform for quantitative evaluation of the folding activity of wildtype and mutant BAM complexes.

**Significance:** The folding of outer membrane proteins (OMPs) in Gram-negative bacteria requires the essential β-barrel assembly machine (BAM). By developing a fluorescent OMP folding reporter, we have unlocked insight into BAM activity *in vitro*, opening the door for rigorous evaluation of BAM mutants and putative inhibitors. We also discovered that, contrary to current models, the BamA POTRA1-3 domains do not contribute significantly to catalysis, despite being essential for bacterial growth. We propose that, consistent with biochemical, structural, and live cell imaging, the *in vivo* role of BamA POTRAs1-3 is to support a connection with the Sec translocon to form a transperiplasmic bridge, as well as provide a docking site for the chaperone SurA for substrate delivery.

## Introduction

Gram-negative bacteria comprise a clinically important subset of bacteria that possess a unique cell envelope; they have two membranes, which are separated by the cell wall-containing periplasm (1). The outer membrane serves a vital function for these bacteria by acting as a selectively permeable barrier, protecting the cell from antibiotics and lipophilic agents (2). Additionally, β-barrel transmembrane outer membrane proteins (OMPs) facilitate important cellular processes such as nutrient transport and drug efflux (3). While the outer membrane presents a roadblock for antibiotic use, an effective antimicrobial strategy might lie in targeting key components of OMP biogenesis (4).

OMPs are ribosomally synthesized in the cytosol then exported to the periplasm by the SEC translocon of the inner membrane (5–7). To prevent aggregation, OMPs can be bound by periplasmic chaperones SurA, Skp, and DegP (8–14). In the currently prevailing model, OMPs are then shuttled by SurA to the β-barrel assembly machine (BAM), which folds and inserts the OMP into the outer membrane (15, 16). BAM function is vital for Gram-negative survival (16, 17).

The prototypical *E. coli* BAM complex consists of the β-barrel OMP BamA and the lipoproteins BamB, C, D, and E (16–19). Cell growth and membrane permeability assays have revealed important insight into the effects of BAM component deletion. BamA and BamD are both conserved in Gram-negatives and essential components as depletion of either is lethal (16, 17). Individual deletion of BamB, BamC, or BamE is tolerated, suggesting these play less vital roles in OMP biogenesis (16, 18, 20). BamA has two structurally distinct domains: the transmembrane β-barrel domain, and the soluble periplasmic polypeptide transport-associated (POTRA) domains, containing 5 repeats in *E. coli* (21–25). BamCDE bind BamA through POTRA 5 and BamB binds BamA through POTRA 3 (18, 24, 26–28). The POTRAs and BamB have also been identified as SurA docking sites (29–32). Intriguingly, POTRA domains 3-5 are essential while deletion of POTRAs 1-2 is tolerated in *E. coli* (24). Accordingly, the POTRA domains have been proposed to bind substrate OMP and may facilitate the formation of β-strands in the OMP substrate (22, 24, 33, 34). However, it is still unclear how these growth phenotypes are related to the OMP folding mechanism of BAM, due, at least in part, to limitations in the available *in vitro* folding assays.

The BAM complex activity can be studied with *in vitro* OMP folding assays, which typically rely on a characteristic “heat modifiability” of OMPs (35) that allows the separation and quantification of folded and unfolded OMP species with SDS-polyacrylamide gel electrophoresis (SDS-PAGE). While accessible, the assay is discontinuous, requires effective quenching and substantial post-processing, and is difficult to implement for fast reactions. An alternative assay uses the OMP protease OmpT as a BAM substrate. As OmpT folds, it cleaves a quenched fluorescent peptide resulting in fluorescence increase (36–41). However, it is difficult to extract the BAM kinetic parameters from this assay.

Here, we report the development of a new platform to quantitatively measure the activity of the BAM complex *in vitro*. We first demonstrate that a fluorescently labeled OMP increases fluorescence when folded into membranes, allowing for folding rates to be quantitatively tracked continuously with high sensitivity and dynamic range. We then use this fluorescent OMP substrate reporter under single-turnover conditions to report kinetic parameters for wildtype BAM complex as well as mutants with deletions of POTRA domains and lipoproteins yielding new insight into BAM function. The results highlight the potential for this new tool in understanding the molecular mechanism of the BAM complex as well as the evaluation of inhibitors.

## Results

### Development of a Novel OMP Folding Reporter

The folding status of OMPs is often evaluated using SDS-PAGE, taking advantage of their characteristic “heat modifiability” (35). In this approach, when OMPs are solubilized and boiled, they denature resulting in a single denatured band in the gels. However, SDS solubilization without boiling does not denature folded OMPs that run with distinct mobility in SDS-PAGE and allows quantification of the folded fraction. We sought to develop a fluorescent reporter to evaluate OMP folding without the need for SDS-PAGE to separate folded and unfolded species. We settled on the transmembrane (TM) β-barrel of OmpA (amino acids 22-197, tOmpA), a construct that has been previously shown to spontaneously fold into liposomes composed of short chain Phosphatidyl-Choline (PC) lipids (12, 42–50). A cysteine was introduced at position three, just outside the TM region, and was labeled with Alexa Fluor 488-maleimide (herein referred to as tOmpA-A488). A folding assay was initiated by mixing of urea-denatured tOmpA-A488 with PC 10:0 liposomes followed by monitoring of fluorescence at 516 nm. We observed a progressive, saturable increase in fluorescence over time (Figure 1B, black dots). Samples were also taken from the folding reaction at different time points and evaluated for folding using the standard SDS-PAGE based “heat modifiability” assay (Figure 1A). There was an excellent correlation between the rate of fluorescence increase and the increase in intensity of the folded tOmpA-A488 band over time (Figure 1B, red diamonds). The change in fluorescence over time was best fit to a double exponential, consistent with previous reports of OMP folding in these lipids (44, 50, 51), which yielded a fast folding rate constant, k_fast_ of 8.5 ± 0.4 min^−1^ (mean ± STD from two independent reactions) (Figure 1I). As a control, mixing of urea-denatured tOmpA-A488 with buffer without liposomes resulted in negligible fluorescence changes and no folding of tOmpA-A488 (Figure 1C).

**Figure 1.**
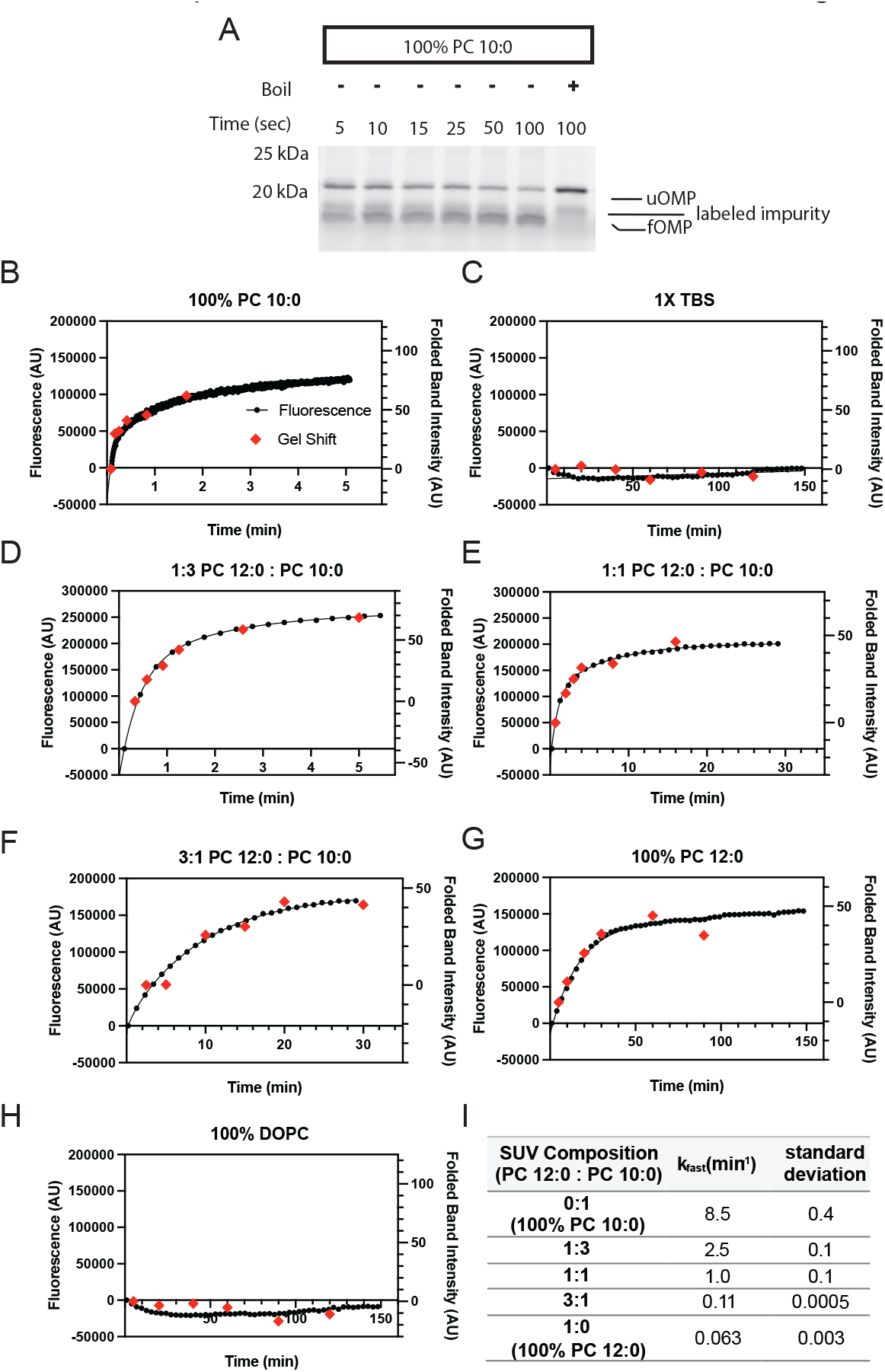
tOmpA-A488 folding monitored by gel-shift and fluorescence. (A) Representative SDS-PAGE heat modifiability assay for tOmpA-A488 folding into PC 10:0 liposomes. Indicated are unfolded (uOMP) and folded (fOMP) tOmpA-A488 and a labeled impurity present in all conditions. (B-H) Representative bulk fluorescence vs time (black dots) of tOmpA-A488 in solutions of mixed PC liposomes (B, D-G), buffer (C), or DOPC liposomes (H), overlaid with data from folding quantification from SDS-PAGE (red diamonds). (I) All fluorescence data were fitted to double exponential functions, except the 1:3 PC10 : PC 12:0 condition, which was fitted to a single exponential, and the fast rate constant is reported as the average of two independent replicates ± STD.

The phospholipid packing order in liposomes increases with the length of the phospholipid acyl chains and decreases the rate of spontaneous OMP folding (51, 52). We thus tested the ability of tOmpA-A488 to report the rate of folding in liposomes with increasing ratios of PC 12:0/PC 10:0 phospholipids. As expected, increased PC 12:0 lipid fraction resulted in decreased rates of tOmpA-A488 fluorescence (Figure 1D-G, black dots). In all cases, fluorescence change showed excellent correlation with the increase in intensity of the folded tOmpA-A488 band over time in the standard “heat modifiability” assay (Figure 1D-G, red diamonds). Double exponential fits yielded k_fast_ values of 2.5 ± 0.1, 1.0 ± 0.1, 0.11 ± 0.0005, 0.063 ± 0.003 min^−1^ (mean ± STD from two independent reactions) for PC 12:0-to-PC 10:0 ratios of 1:3, 1:1, 3:1, and 1:0 (100% PC 12:0), respectively (Figure 1I). Furthermore, we tested folding of tOmpA-A488 into liposomes composed of standard-length PC 18:1(Δ9-cis) phospholipids (DOPC), which do not allow spontaneous OMP folding (51). As expected, negligible fluorescence changes and no folding of tOmpA-A488 were observed in the time frame of the experiment in these liposomes (Figure 1H). Taken together, these results indicate that tOmpA-A488 is a suitable reporter to evaluate folding by direct monitoring of fluorescence changes at 516 nm.

### Single Turnover Kinetics of the BAM Complex Reveals the Rate of Folding Catalysis and Estimates of Substrate Affinity

We next used the tOmpA-A488 reporter to quantify folding catalysis by BAM. Wildtype BAM complex (BamABCDE) was reconstituted into liposomes composed of *E. coli* polar lipids (EPL) using a previously described dialysis method (38). Standard folding assays were carried out by mixing urea-denatured tOmpA-A488 with the BamABCDE proteoliposomes and the periplasmic chaperone SurA (38, 46, 49). Due to OMP aggregation propensity and a potential for limited capacity of OMP folding within highly packed proteoliposomes, we chose to utilize single-turnover conditions to probe BAM catalysis where [tOmpA-A488] << [BAM]. To verify such conditions, we tested tOmpA-A488 concentrations ranging from 2 to 50 nM while keeping the BAM concentration at 0.75 µM. We observed exponential increases in fluorescence that scaled with the tOmpA-A488 concentration but resulted in indistinguishable exponential folding rates confirming single-turnover conditions (Figure S1). Next, we carried out folding assays at 10 nM tOmpA-A488 and increasing concentrations of BamABCDE proteoliposomes in the µM range that resulted in exponential fluorescence increases that saturated at rates proportional to the concentration of BAM (Figure 2A). Reaction endpoints subjected to SDS-PAGE under heat modifiability conditions confirmed that the fluorescence increases correlated with tOmpA-A488 folding (Figure 2A, inset). The single exponential folding rate constants obtained from the curves in Figure 2A were plotted against the concentration of BAM to yield a hyperbolic curve that was fitted to an x-axis “offset” square hyperbola (Eq. 1, see methods). The asymptote reflects the intrinsic rate of BAM catalysis (k_fold_) observable at full substrate saturation, while the concentration of BAM at half the k_fold_ represents the constant value, K_0.5_, composed of the substrate association (k_1_) and dissociation (k_-1_) rate constants, and k_fold_ (Scheme 1, Eq. 2). We report that wildtype BamABCDE complex has a k_fold_ of 0.78 ± 0.15 min^−1^ and K_0.5_ of 3.1 ± 1.1 µM (mean ± STD from three independent BAM reconstitutions). Assuming the 0.78 ± 0.15 min^−1^ k_fold_ is slow compared to the substrate dissociation constant k_-1_, the K_0.5_ can serve as an approximation of the substrate dissociation constant (K_d_). These parameters were highly reproducible across separate BAM protein preparations and reconstitutions. Importantly, we saw no folding of tOmpA-A488 into empty EPL liposomes, confirming that the folding we observed is BAM-catalyzed (Figure S2). Furthermore, the k_fold_ we observe is predicted to support normal doubling times in *E. coli* (53).

**Figure 2.**
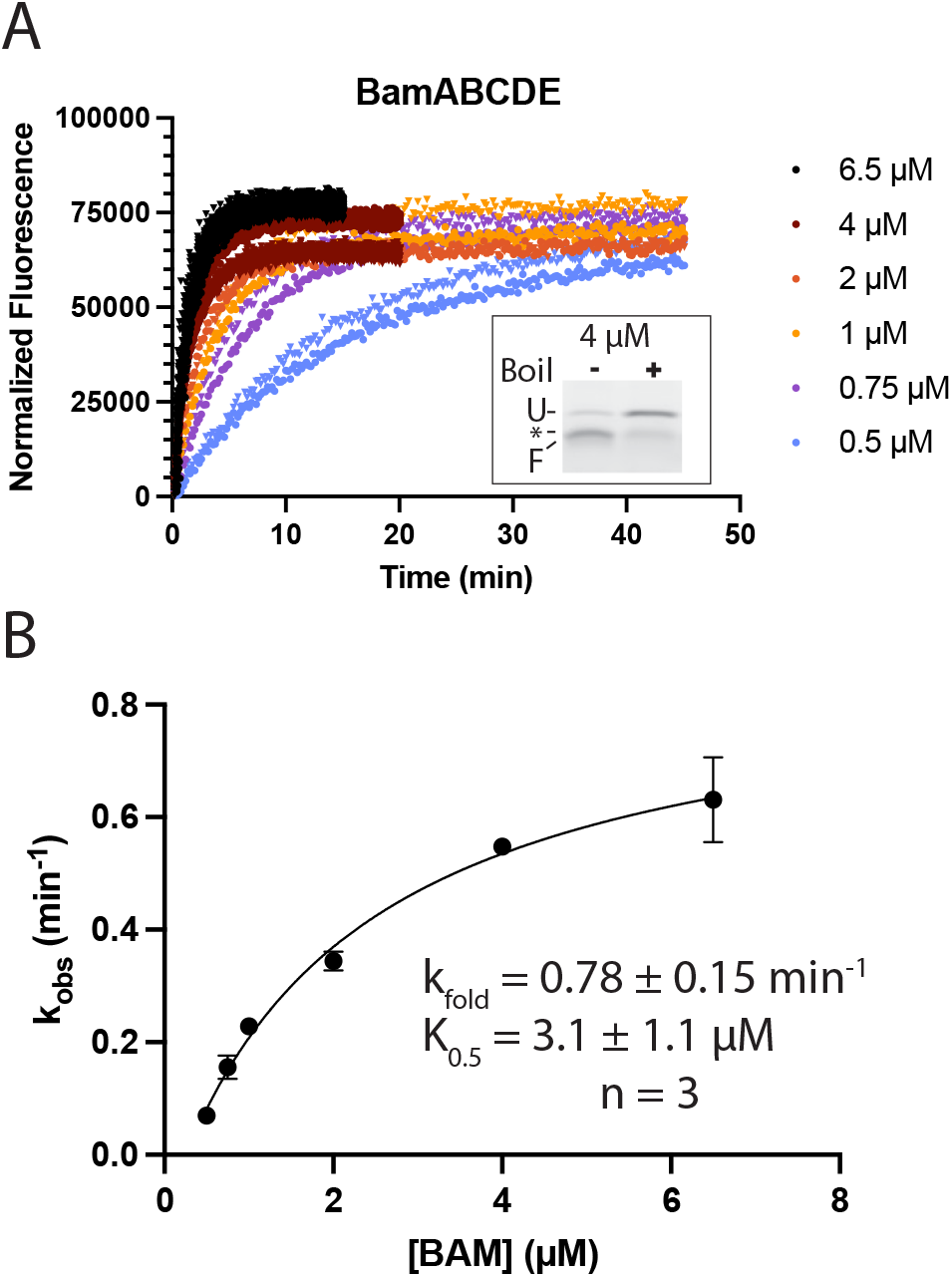
Single-turnover kinetics of wildtype BamABCDE complex. (A) Representative fluorescence vs time data from a BAM activity assay. Background corrected data were fitted to a single exponential function and normalized to the Y-intercept value. Technical replicates for given concentrations are shown as triangles of the same color. Inset: representative heat modifiability assay for tOmpA-A488 with 4 µM BAM after 20 minutes. Indicated are unfolded (U) and folded (F) tOmpA-A488, and a fluorescent contaminant (asterisk). (B) Single exponential rate constants from (A) were fitted to an “offset” hyperbolic curve (Eq. 1). Data points are the mean of two technical replicates, with SD error bars. Inset: fit parameters for three independent biological replicates, reported as the mean ± SD.

### The BAM Complex is Active Without the First Three POTRA Domains *in vitro*

Having reported robust activity of wildtype BamABCDE, we next tested what role the POTRA domains play in OMP folding. We constructed a BAM mutant with a deletion of BamA POTRAs 1-3 (BamA ΔP1-3). Previous growth complementation assays show that deletion of POTRAs 1 and 2 are tolerated in *E. coli* but lead to growth defects whereas deletion of POTRA3 is lethal (24, 54). Therefore, we hypothesized that deleting the first three POTRA domains would be deleterious to activity. SDS-PAGE confirmed purification of the expected BamA truncation, with minimal contamination from wildtype endogenous BamA (Figure S3). As expected, the ΔP1-3 mutant did not contain BamB, which interacts primarily with BamA POTRA3 (27), resulting in a BamA(ΔP1-3)CDE complex. Single-turnover activity assays revealed that the BamA(ΔP1-3)CDE complex was active, and substrate saturation was approached with the concentrations of BAM tested (Figure 3A). The rate constants were plotted as a function of BAM concentration and fitted to a square hyperbola as described above (Figure 3B). Remarkably, the analysis revealed the ΔP1-3 mutant showed only a modest decrease in activity, with a k_fold_ of 0.41 ± 0.11 min^−1^ (mean ± STD from three independent BAM reconstitutions). Strikingly, the K_0.5_ of the BamA ΔP1-3 mutant was no different than WT BAM, suggesting that removal of the three terminal POTRA domains, as well as BamB, has little effect on the substrate binding affinity. Similar results were obtained for BamA(ΔP1)BCDE and BamA(ΔP1-2)BCDE complexes (Figure S4A-D), further illustrating the limited role of POTRAs 1-3 in substrate binding and BAM catalysis under our experimental conditions.

**Figure 3.**
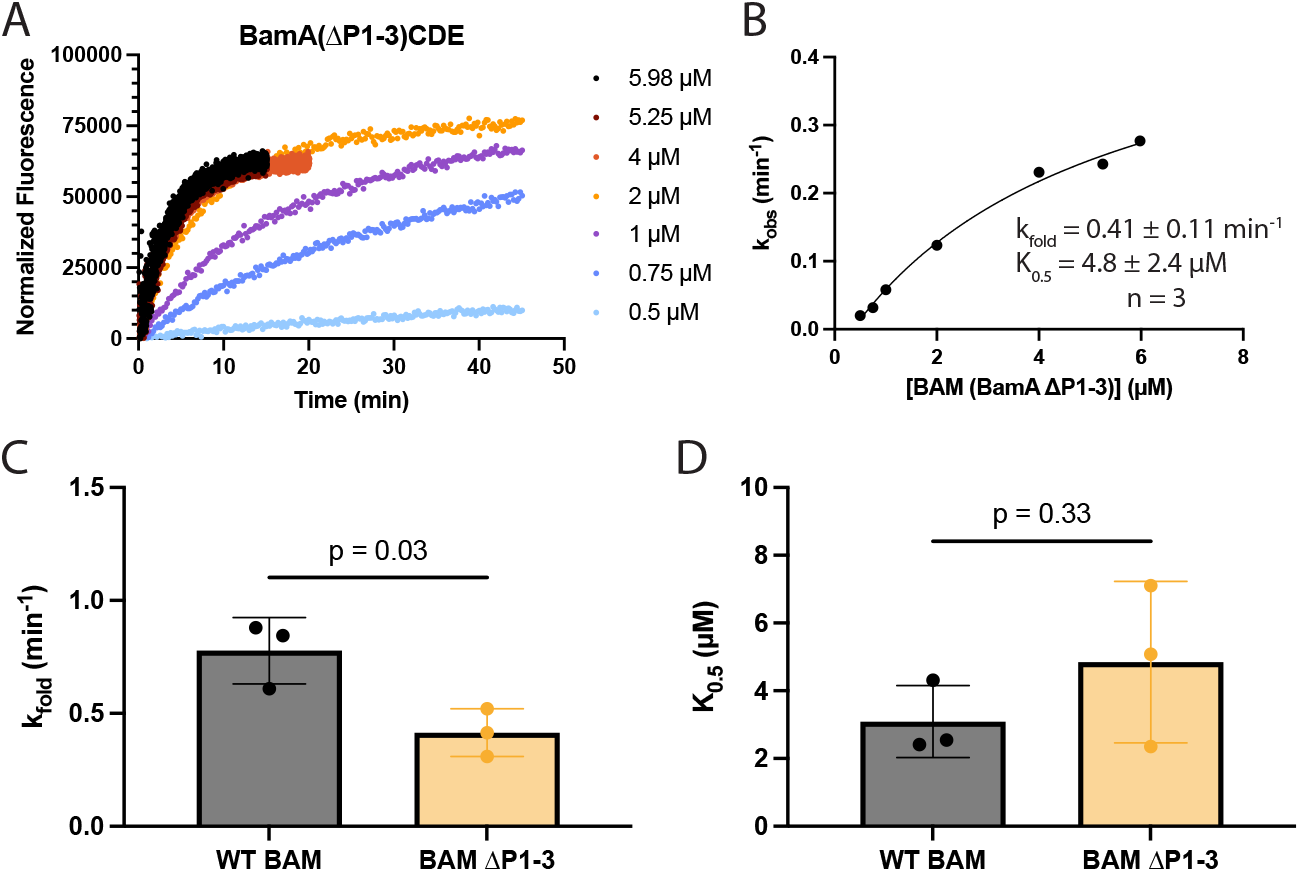
Single-turnover kinetics reveal that BAM is catalytically active without POTRA domains 1-3. (A) Representative fluorescence vs time data from single-turnover activity assay of the BamA(ΔP1-3)CDE mutant. (B) Data from (A) were fitted to Eq. 1. Inset: fit parameters for three biological replicates, reported as the mean ± SD. (C-D) Calculated k_fold_ and K_0.5_ parameters, respectively for WT and the ΔP1-3 BAM mutant. An unpaired Welch’s t-test indicated a p-value of 0.03 and 0.33, respectively between WT BAM (n=3) and BAM ΔP1-3 (n=3).

### BamA by itself is Defective at Catalyzing OMP Folding in *E. coli* Membranes and the E470K Mutation does not Rescue Activity

As deletion of BamB and three POTRA domains had little effect on tOmpA-A488 folding activity, we next tested the effect of removing all BAM lipoproteins. To probe the catalytic capability of BamA by itself, we expressed a His-tagged BamA that complements BamA depletion in *E. coli* indicating that the tag is well tolerated (34). Purification of this protein yielded BamA with only trace amounts of co-purified endogenously expressed BamB, BamC, and BamD (Figure S5). Single-turnover kinetic assays with BamA in EPL liposomes show minimal changes in tOmpA-A488 fluorescence indicative of poor-to-no folding (Figure 4A), consistent across three independent biological replicates. SDS-PAGE analysis confirmed that the reconstituted BamA was folded (Figure S5), however tOmpA-A488 was not folded by BamA, while it is nearly completely folded under similar conditions with wildtype BamABCDE (Figure 4D). These results indicate that at concentrations close to saturation for WT BamABCDE complex, BamA by itself is essentially incapable of catalyzing OMP folding in *E. coli* lipid membranes. Silhavy and colleagues have recently reported a gain-of-function mutation in BamA (55). Cell-based assays suggest that BamA(E470K)BCDE complex is phenotypically indistinguishable from wildtype BamABCDE (56). However, cells expressing BamA(E470K) can survive depletion of the essential lipoprotein BamD in a BamBCE knockout strain, indicating that BamA(E470K) by itself is able to support cell growth (55, 57). Therefore, we hypothesized that BamA(E470K) may rescue the in vitro activity of BamA alone. BamA(E470K) from inclusion bodies to limit co-purification of trace amounts of lipoproteins, which may contribute background levels of activity. The purified protein was then solubilized in tris-buffered urea, refolded in detergent, and subjected to size exclusion chromatography (see methods). We also expressed BamA(E470K) with its signal sequence and purified it from *E. coli* membranes. Neither refolded nor membrane-extracted BamA(E470K) demonstrated heat modifiability, consistent with previous reports (Figure S6A) (56). However, the BamA(E470K) proteins eluted similarly to membrane-extracted BamA and circular dichroism demonstrated similar secondary structure as wildtype BamA (Figure S6B). Thus, our refolded and membrane-extracted samples of BamA(E470K) likely adopt the BamA native fold. Interestingly, single-turnover kinetics measurements of the refolded and membrane-extracted BamA(E470K) mutant alone showed little-to-no tOmpA-A488 folding activity (Figure 4B-C). Furthermore, a purified BamA(E470K)BCDE complex showed activity similar to wildtype BamABCDE complex in our single-turnover experiments (Figure S7), consistent with the similar phenotypes observed in cell-based assays (56). Together, these results indicate that the BamA(E470K) mutation does not dramatically enhance the BAM complex k_fold_, nor does it bring the K_0.5_ for BamA by itself to values similar to the wildtype BamABCDE complex.

**Figure 4.**
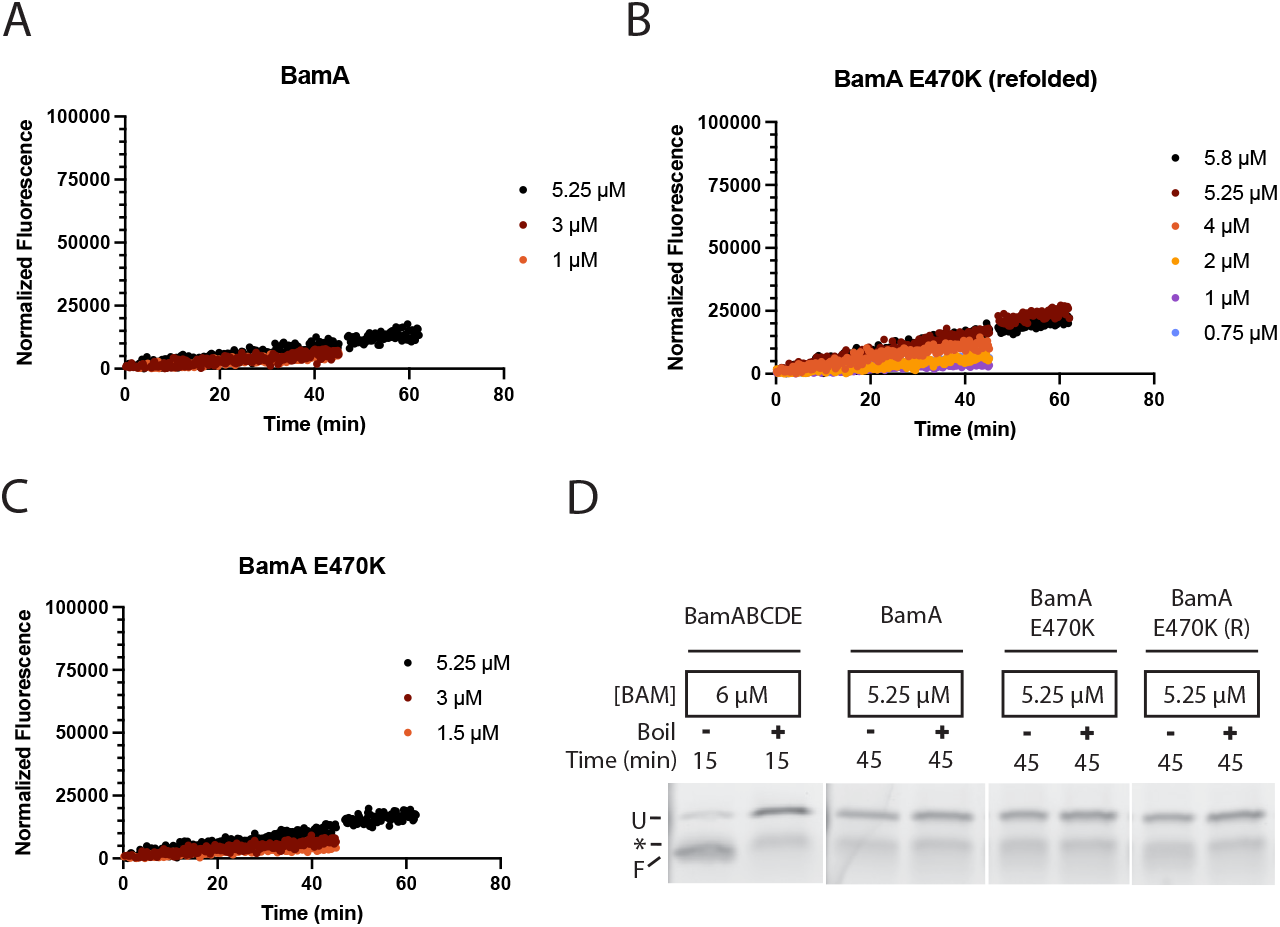
BamA alone is defective in activity. (A) Representative fluorescence vs time data from a single-turnover activity assay of BamA alone. (B-C) Fluorescence vs time data from single-turnover activity assay of refolded BamA (E470K) and membrane-extracted BamA (E470K), respectively. (D) Fluorescent images of SDS-PAGE of tOmpA-A488 folding reactions with BamABCDE complex, wildtype BamA alone, BamA(E470K) alone, and refolded BamA(E470K) (R).

## Discussion

The most frequently used approach to evaluate BAM complex folding of OMPs relies on their “heat modifiability” and the use of SDS-PAGE to separate folded and unfolded species (35). While accessible and in principle applicable to any OMP substrate, the approach is discontinuous, affords limited sampling of fast kinetics, requires effective folding quenching and substantial postprocessing (SDS-PAGE and quantification). A commonly used alternative to measure BAM activity is based on folding OmpT, an OMP with protease activity which cleaves a quenched fluorescent peptide (36–41). The resulting increase in fluorescence over time can be monitored continuously with high sensitivity in this form of coupled assay. However, extracting the kinetic parameters for the BAM catalyzed reaction is difficult as the first reaction (OmpT folding) produces the catalyst (OmpT) for the measured reaction (peptide cleavage) resulting in complex kinetic behavior. To address these challenges, we developed a novel fluorescent OMP folding reporter based on the β-barrel transmembrane domain of OmpA labeled with Alexa 488 (tOmpA-A488). We demonstrate that tOmpA-A488 change in fluorescence robustly tracks its spontaneous folding into short-chain PC liposomes (evaluated by the standard SDS-PAGE assay) under multiple conditions where the intrinsic folding rates varied by over two orders of magnitude while showing no fluorescence change when no folding occurs (Fig. 1). Therefore, we attribute tOmpA-A488 change in fluorescence to an increase in its fluorescence quantum yield when the protein adopts its folded structure in the membrane.

A central challenge in measuring the folding activity of the BAM complex is the proclivity of its unfolded OMP substrates to aggregate or misfold, which compete with productive folding (58). This has hindered previous attempts to study BAM kinetics using steady state approaches requiring saturating concentrations of substrate (45). To overcome this challenge, we took advantage of the high sensitivity of the tOmpA-A488 reporter to set up single-turnover reaction conditions with nanomolar concentrations of unfolded substrate and micromolar concentrations of BAM complex to minimize the aggregation problem. We observed saturable increase in fluorescence that fit to single exponential rates and scaled hyperbolically with the concentration of BAM complex (Fig. 2). Fitting of these data to an offset hyperbolic equation (Eq. 1) allows the estimation of the BAM complex catalytic folding constant k_fold_ of 0.78 ± 0.15 min^−1^ (mean ± STD from three independent BAM reconstitutions). This value may seem very slow compared with ~600 min^−1^ typical of enzymes that catalyze chemical reactions (59). However, we interpret the k_fold_ as the overall rate constant for the entire folding process catalyzed by the BAM complex as analysis of the reaction endpoints by SDS-PAGE indicated heat-modifiable, folded species of tOmpA-A488 (Fig. 2A). According to the current mechanistic model, OMP folding involves sequential threading of substrate β-hairpins in the BamA barrel gate to form hybrid barrels, followed by budding of the completed barrels into the membrane (60–63). This process is likely to be significantly slower than enzyme catalyzed chemical conversions. In fact, our reported k_fold_ is approximately 10-fold faster than previous estimates (45), and is in excellent agreement with the BAM k_fold_ of ~0.6 min^−1^ estimated using deterministic and stochastic models to simulate rates of OMP folding needed to sustain normal doubling times in *E. coli* (53). Therefore, the reported k_fold_ is consistent with current estimates of what is required for normal physiological conditions.

In addition to the k_fold_, analysis of the single-turnover kinetic data also yields K_0.5_, which represents the combination of the k_fold_ and the substrate binding and unbinding rate constants (K_0.5_ = (k_-1_+k_fold_)/k_1_). Assuming that substrate unbinding is much faster than the folding (k_-1_>>k_fold_), a conservative assumption given the slow k_fold_ of 0.78 ± 0.15 min^−1^, the K_0.5_ approximates the substrate K_d_. We observe a K_0.5_ of 3.1 ± 1.1 µM (mean ± STD from three independent BAM reconstitutions) for wildtype BamABCDE. As the BAM proteoliposome reconstitution is the most challenging aspect to replicate from assay to assay, it likely introduces the most variability. Different reconstitutions may result in variable yields of active enzyme, namely through differential complex orientation in the liposomes (i.e. POTRA domains facing outwards versus inwards) or entrapment of liposomes inside other liposomes (i.e. multilamellar vesicles). Whereas this variability is not expected to affect the k_fold_, an intrinsic property of the enzyme, different reconstitution efficiencies affect the estimate of K_0.5_, as this depends on the concentration of active enzyme. This is evidenced by a somewhat larger standard deviation of the K_0.5_ estimates compared to the k_fold_ value. Nevertheless, the K_0.5_ of 3.1 ± 1.1 µM for BAMABCDE is comparable to binding affinities of proteins in the OMP biogenesis pathway, where the reported K_d_ of SurA to tOmpA and SurA to BAM are 1.8 and 2.6 µM, respectively (12, 31). Taken together, this indicates that our observed kinetic parameters are physiologically reasonable. Furthermore, the experiments establish the use of the tOmpA-A488 folding reporter in combination with single-turnover kinetics as a robust platform to quantitatively test the activity of BAM.

To understand the effect of BamBCDE lipoproteins on the complex activity, we tested the activity of BamA by itself in liposomes composed of *E. coli* lipids. Several studies have previously demonstrated that BamA by itself can accelerate OMP folding into short-chain PC (12) and mixed PC/PE liposomes (45, 48, 64), suggesting BamA is responsible for driving the catalytic folding of OMPs. Notably, these liposomes with non-physiological phospholipids allow OMP folding independent of BAM (45, 51, 64), which complicates extrapolation of folding rates to physiological conditions. Conversely, liposomes composed of *E. coli* polar lipids do not allow spontaneous insertion of OMPs (51). Therefore, we reconstituted BamA into liposomes composed of *E. coli* lipids such that any OMP folding activity could be ascribed to BamA. Unlike the wildtype BamABCDE complex, BamA by itself was unable to efficiently fold tOmpA-A488 in our single-turnover experiments (Fig. 4). This is consistent with previous reports of poor folding of tOmpA by BamA in *E. coli* proteoliposomes (less than 20% after two hours) (49) and is consistent with the lethality of BamD deletion in cells (17). As the gain-of-function mutant BamA(E470K) has been reported to bypass the essential requirement for BamD in *E. coli* (55, 57), we hypothesized that the BamA(E470K) mutant might be more active than wildtype BamA in our assay. However, when reconstituted into *E. coli* phospholipid liposomes, BamA(E470K) by itself was just as defective as wildtype BamA at folding tOmpA-A488. Furthermore, BamA(E470K)BCDE complex did not show enhanced activity compared to wildtype BamABCDE complex. Combined, these results indicate that the E470K mutation does not dramatically improve the BAM complex k_fold_ nor does it appear to increase productive substrate binding as previously proposed (65). Therefore, the mechanism by which the BamA E470K mutation bypasses the requirement for BamD in *E. coli* cells remains elusive.

Data from multiple labs suggested that BamA POTRA domains play a role in substrate binding (22, 24, 33). Specifically, because β-augmentation, or β-strand templating, was frequently observed for POTRA3 with neighboring POTRA domains in crystal lattices, it was hypothesized that one function of the POTRA domains might be to bind substrate OMPs and nucleate the formation of β-strands in a similar manner (22, 24). Furthermore, we previously identified that conformational flexibility between the POTRAs 2 and 3 was important for *E. coli* cell growth (34). This led to the hypothesis that conformational ratcheting between POTRA domains may facilitate formation of β-hairpins between OMP β-strands bound to neighboring POTRA domains, thereby helping initial folding of OMPs. Moreover, the lethality of a POTRA 3 deletion led to the hypothesis that POTRA3 may specifically serve an important role in OMP binding aside from its function to bind the non-essential lipoprotein BamB (24). To probe these hypotheses, we constructed BAM complexes with increasing severity of POTRA deletions: BamA(ΔP1)BCDE, BamA(ΔP1-2)BCDE, and BamA(ΔP1-3)CDE. Based on the accepted model in the field that POTRA domains are important for binding OMPs, we predicted the POTRA deletions would have progressively bigger impacts on substrate binding. Additionally, because of the lethality in cells (24), the POTRA1-3 deletion, which also removes BamB from the BAM complex, was expected to have a severe impact on activity. However, single-turnover assays of the BamA(ΔP1-3)CDE complex revealed a very modest impact on k_fold_ (0.41 ± 0.11 versus 0.78 ± 0.15 min^−1^, p=0.03). The small reduction in k_fold_ is unlikely to be responsible for lethality because a 5-10-fold reduction in BamA expression is well tolerated in *E. coli* (66). Furthermore, control experiments with POTRA 1 and POTRA1-2 deletion mutants also yielded k_fold_ values similar to those of the POTRA1-3 deletion mutant and those mutants are tolerated in cell growth assays (24, 54). The observed k_fold_ is also well within the range (between 0.18 and 3.6 min^−1^) predicted in OMP folding simulations to support normal cell growth (53). The data indicate that the first 3 POTRA domains do not play a substantial role in folding catalysis and therefore does not support the POTRA ratcheting model of forming OMP β-hairpins. We also found no significant difference in K_0.5_, or apparent substrate binding affinity, between the complexes with POTRA1-3 deletions and wildtype BamABCDE (Fig. 3). Taken together, the data indicate that the first three domains do not significantly contribute to OMP binding under our single-turnover conditions. It is possible that the lethality of the POTRA3 deletion is due to a role in substrate turnover that is not captured in our assay. However, we propose a different alternative.

The prevailing model for OMP biogenesis calls for the chaperone SurA to capture nascent OMPs as they emerge from the Sec translocon in the inner membrane shuttling them to the BAM complex (32). However, SurA is not an essential protein in *E. coli* (32, 67) and while other periplasmic chaperones such as Skp, DegP and FkpA may be involved in aspects of OMP biogenesis, they do not seem to play a role in substrate delivery to the BAM complex. This suggests the existence of an alternative pathway for substrate delivery. There is growing evidence that the Sec translocon machinery directly interacts with the BAM complex to form a transperiplasmic bridge allowing direct transfer of nascent OMPs to the BAM complex for insertion in the outer membrane (54, 68–71). Using single molecule analysis in live cells, we recently showed that diffusion of the Sec holotranslocon is unusually slow for an inner membrane protein, approaching that of the BAM complex in the outer membrane (54). However, Sec diffusion is significantly increased when the BAM complex has a POTRA1 or POTRA1-2 deletion suggesting a Sec-BAM interaction. Consistent with these experiments, we propose that the main role of POTRA1-3 domains is to support a connection between the BAM complex and the Sec holotranslocon allowing direct transfer of nascent OMPs to the outer membrane. Thus, we suggest a model where OMPs transfer to BAM via parallel pathways, one involving direct transfer through a Sec-BAM bridge, and another dependent on SurA to shuttle nascent OMPS that fail to engage the bridge back to BAM. In this model, disruption of one pathway leads to outer membrane defects and/or impaired growth, while disruption of both pathways is lethal. Such model is consistent with the observed mutant phenotypes. A SurA deletion is not lethal as it disrupts only one pathway. Similarly, POTRA1 or POTRA1-2 deletions are not lethal as they disrupt the bridge (54), but SurA chaperoning would still occur. Recent cryo-EM data indicates that SurA binding to BAM is mediated by interactions with BamB as well as β-augmentation of POTRA1 by the flexible, unstructured N-terminus of SurA (30, 72). As β-augmentation interactions are sequence promiscuous, we speculate that in the POTRA1 and POTRA1-2 deletions, SurA may be able to dock to BAM via β-augmentation of POTRA2 or 3 with its flexible N-terminus, while retaining the interactions with BamB. However, if POTRAs1-3 are deleted (which also removes BamB from the complex), the Sec-BAM bridge is disrupted, and SurA loses its BAM docking sites disrupting the chaperoning pathway resulting in a lethal phenotype.

A physiological role for a Sec-BAM transperiplasmic bridge in OMP biogenesis is supported by biochemical experiments in independent laboratories (68–71) as well as our live cell microscopy data (54). The *in vitro* single-turnover kinetics experiments presented here are consistent with a role for the first three POTRA domains of BamA in formation of such bridge rather than a role in substrate binding or folding catalysis. On the other hand, some estimates of the distance between the inner and outer membrane may be inconsistent with a direct interaction between Sec and BAM (73). This may suggest that additional periplasmic components important for OMP biogenesis are yet to be discovered.

## Materials and Methods

Experimental procedures for cloning and protein expression and purification are provided in SI Materials and Methods. Plasmids and oligonucleotides used are detailed in Tables S1 and S2.

### Labeling of tOmpA 1 Cys N-term with Alexa Fluor 488

The guanidine-HCl stock of tOmpA 1 Cys N-term was diluted 10-fold into 8 M urea, 20 mM Tris, 1 mM EDTA, 0.05 mM TCEP, pH 7.3, to yield ~100 µM OMP. The protein was reduced with 30 mM TCEP at 21°C for 30 minutes. Protein was acetone precipitated (protocol adapted from Thermo Scientific) to remove excess reducing agent. Pellet was washed with cold 1 mM EDTA pH 6 and redissolved in urea buffer. Alexa Fluor-488 C_5_ maleimide was added to protein at a molar ratio of about 5:1 dye to Cys and incubated for 1 hour at 21°C, after which 2 mM N-ethylmaleimide (NEM) was added to block unreacted cysteines and incubated for 10 minutes. Two rounds of acetone precipitation were performed to remove unreacted dye. The pellet was washed with 100% ethanol, dried with nitrogen gas, and resuspended in 8 M urea, 25 mM glycylglycine pH 8. A NanoDrop was used to check protein and dye concentration with appropriate extinction coefficients. Labeled protein was aliquoted into small portions, snap froze with liquid nitrogen and placed at −70°C until use.

### Folding of tOmpA 1 Cys N-term-A488 into PC Liposomes

Solutions of PC liposomes (prepared as described in SI Materials and Methods), UltraPure BSA (AM2616, Invitrogen), urea buffer (8 M urea, 20 mM Tris pH 8), and 1X TBS + 1 mM EDTA were made to desired concentration of liposomes. tOmpA 1 Cys N-term-A488 was added to liposome solution with continuous mixing to initiate folding reactions. Final reaction concentrations were 0.041-0.8 mM PC lipids, 0.5 mg/mL BSA, 0.8 M urea, 50 nM OMP. Procedures for solution fluorescence measurements and quantification of folding of tOmpA-A488 by SDS-PAGE are detailed in SI Materials and Methods.

### Single Turnover BAM Activity Assays

DDM solubilized BAM was reconstituted into liposomes using a dialysis protocol modified from (38). The resulting proteoliposomes were extruded and final BAM concentration was assayed by BCA assay (a detailed description is provided in SI Materials and Methods). Solutions of BAM proteoliposomes/SurA/1X TBS + 1 mM EDTA buffer were placed in cuvettes on the fluorimeter with stirring. Background fluorescence measurements were taken, as described above, then reaction was initiated by addition of 0.1 µM tOmpA-A488 in 8M urea, 20 mM Tris pH 8 and fluorescence was monitored at appropriate times. Final reaction concentrations were 0.1 µM SurA, 0.01 µM tOmpA-A488, 0.8 M urea, and variable amounts of BAM proteoliposomes (the same proteoliposome reconstitution was used to conserve reconstitution efficiency across a single biological replicate. Processing of the data was performed as described in SI Materials and Methods in Prism 10. Activity curves were constructed by plotting the single exponential folding rate constants (k_obs_) against the concentration of BAM. Activity is modelled after Scheme 1, and activity curves were fitted to an adapted hyperbolic curve described by Eq. 1:

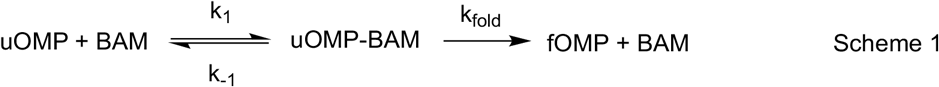

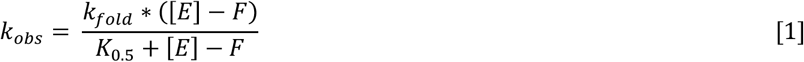

Where [E] represents the BAM concentration, F represents the x-intercept (allows variation in BAM quantification by the BCA assay), and K_0.5_ described by Eq. 2:

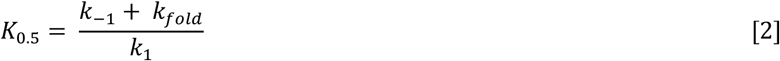

## Acknowledgements

We thank Dr. Harris Bernstein for donating the pJH114 plasmid to our lab. Much thanks to Dr. Cristina Sandoval for the construction and cloning of the tOmpA 1 Cys N-term plasmid and the purification of SurA. We thank the Biochemistry Department Shared Instruments Pool at CU Boulder (RRID: SCR_018986) for use of its general infrastructure (centrifuges, Avestin C3 homogenizer, QM-6 PTI spectrofluorometer, the Tecan Spark Multimode Plate Reader, and the Amersham Typhoon 5 Imager). The Typhoon 5 was funded by NIH S10OD034218. We are immensely grateful for the support provided by SIP staff members Dr. Annette Erbse and Emily Proksch. This work was supported in part by NIH grant 1R01GM127462 to MCS and NIH Signaling and Cellular Regulation T32 Training Grants GM142607 and GM008759 supporting WNB.

