## Supplemental Information for "A fluorescent reporter and single-turnover kinetics reveal new insight into BAM complex function"

### Supplementary Information

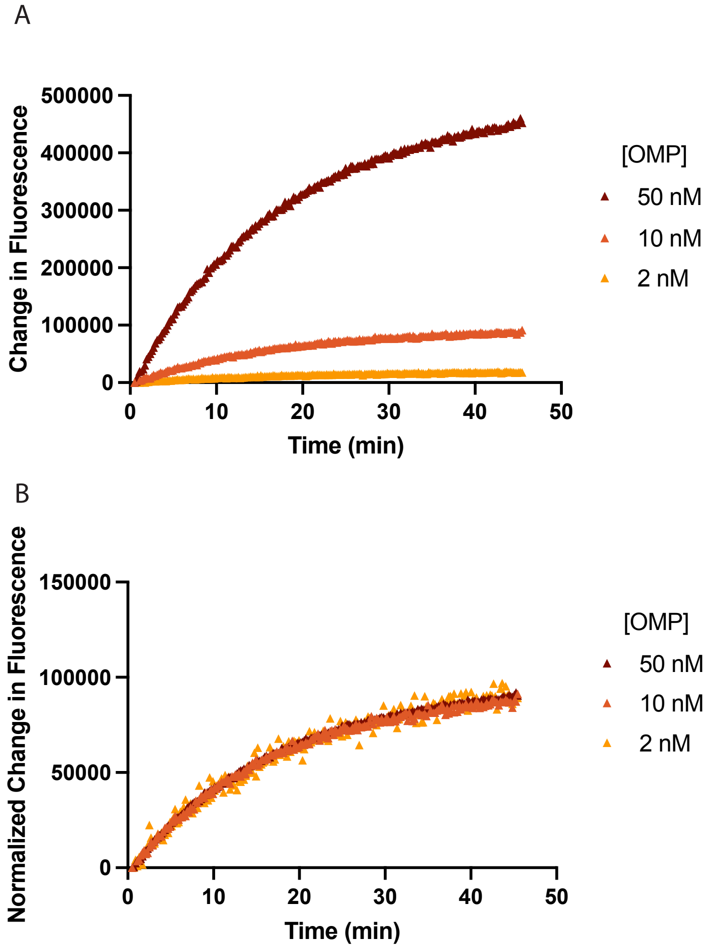


Fig. S1. Effect of changing the tOmpA-A488 concentration under single-turnover activity assay conditions
(A) Fluorescence of tOmpA-A488 folding with 0.75 μM BamABCDE proteoliposomes. (B) Fluorescence from (A) corrected by concentration of tOmpA-A488 normalized to the 10 nM OMP condition. Fluorescence values for the 50 nM condition

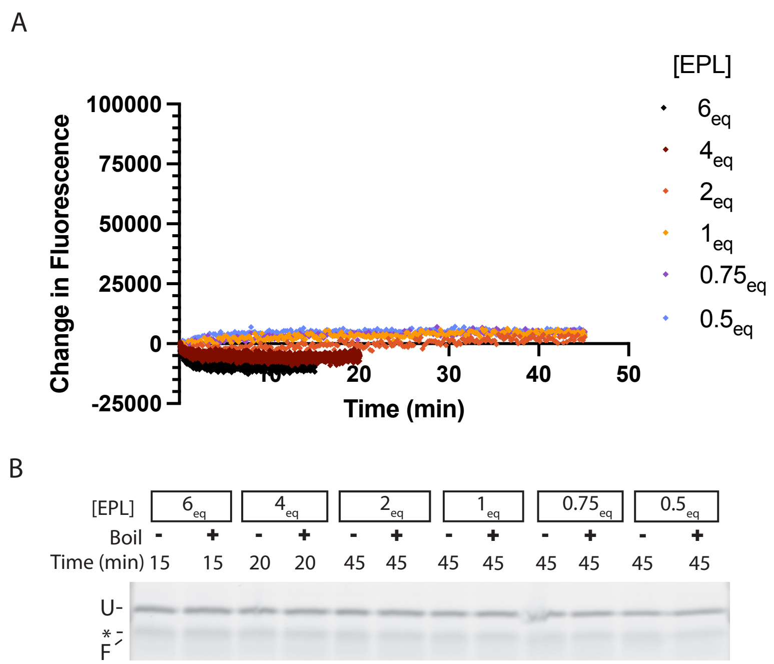


Fig. S2. tOmpA-A488 shows no folding in reconstituted empty EPL liposomes
(A) Fluorescence of tOmpA-A488 in solutions of empty EPL liposomes at roughly equivalent (eq) concentrations of BAM proteoliposomes typically used. (B) Fluorescent image of an SDS-PAGE heat modifiability assay of reactions from (A). Indicated are unfolded (U) and folded (F) tOmpA-A488, and a non-specific fluorescent contaminant (asterisk).

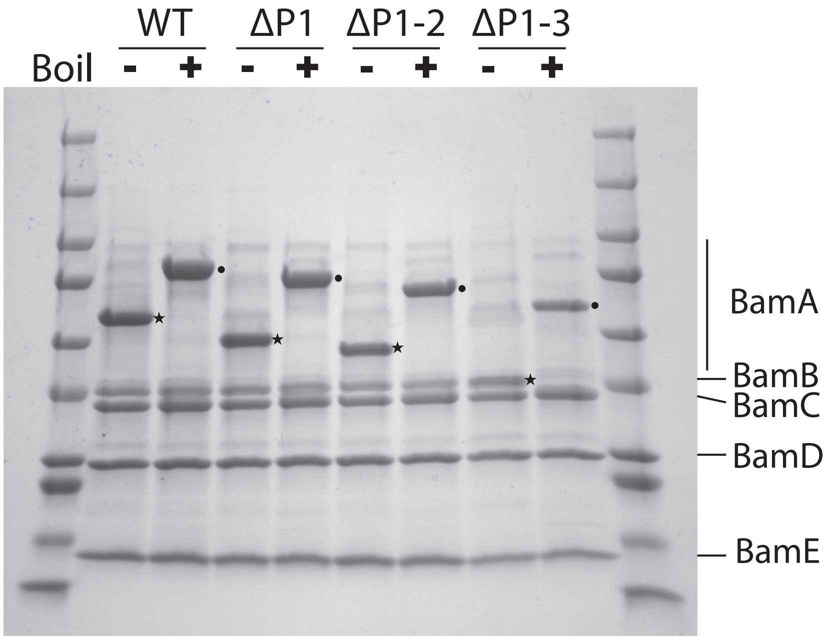


Fig. S3. Purification of BAM POTRA deletion mutants
SDS-PAGE of purified BAM complexes with BamA truncations. Highlighted are the change in molecular weight of the BamA proteins, while BamB, BamC, BamD, and BamE are also present. Note that for the ΔP1-3 mutant, folded BamA runs at a similar molecular weight as BamB. Folded and unfolded BamA bands are indicated with stars and circles, respectively.

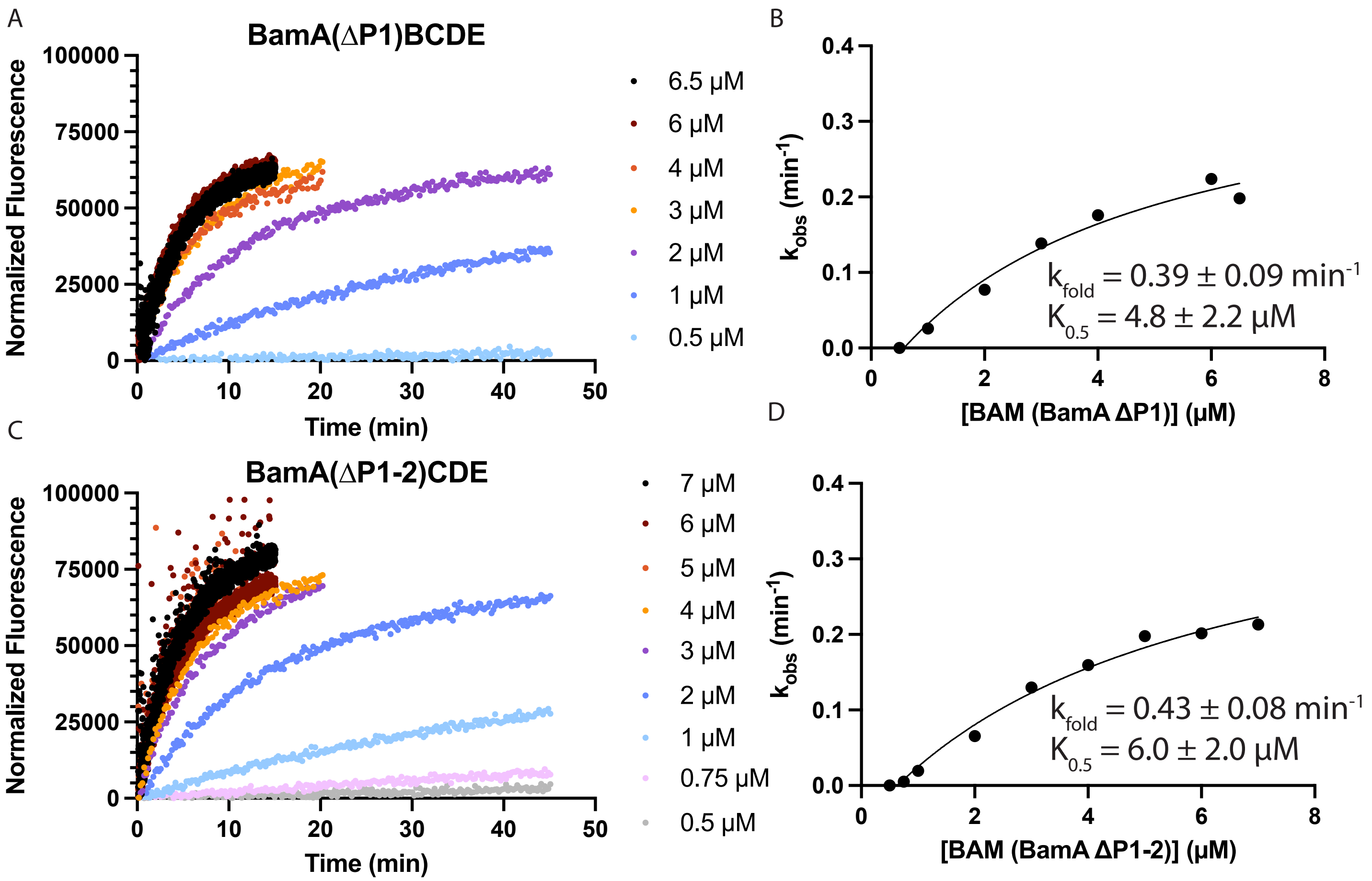


Fig. S4. Single-turnover activity assays of BAM ΔP1, BAM ΔP1-2, and WT BAM without SurA.
(A) Fluorescence data from one independent single-turnover activity assay of the BAM ΔP1 mutant at different concentrations. (B) Single exponential rate constants from (A) were plotted against BAM concentration and fitted to Eq. **Error! Reference source not found.** and represents one biological replicate. Values for $k_{fold}$ and $K_{0.5}$ represent the e stimated value ± SE of the fit. Fluorescence data and fitted rate constants are shown in (C-D) for the BAM ΔP1-2 mutant.

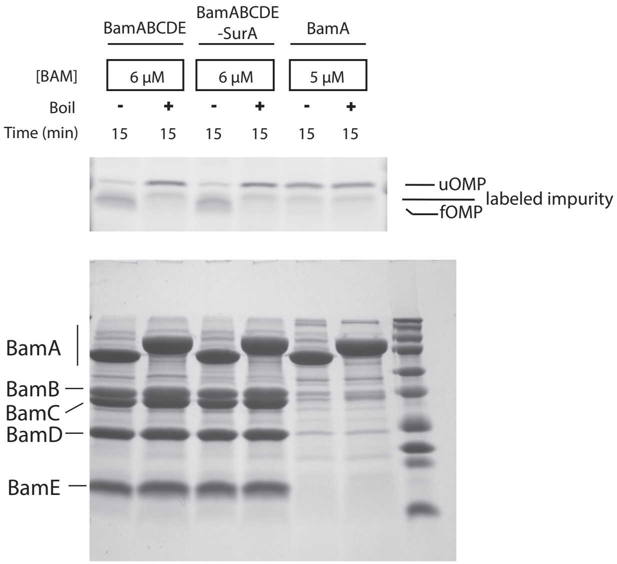


Fig. S5. SDS-PAGE confirms that reconstituted BamA is folded but not active.
(Top) Fluorescent image of an SDS-PAGE heat modifiability assay of reactions with BamABCDE or BamA only after 15 minutes. Indicated are unfolded (uOMP) and folded (fOMP) tOmpA-A488, and a non-specific fluorescent impurity. (Bottom) Coomassie-stained gel from (A). BamA only is folded and has trace amounts of lipoproteins.

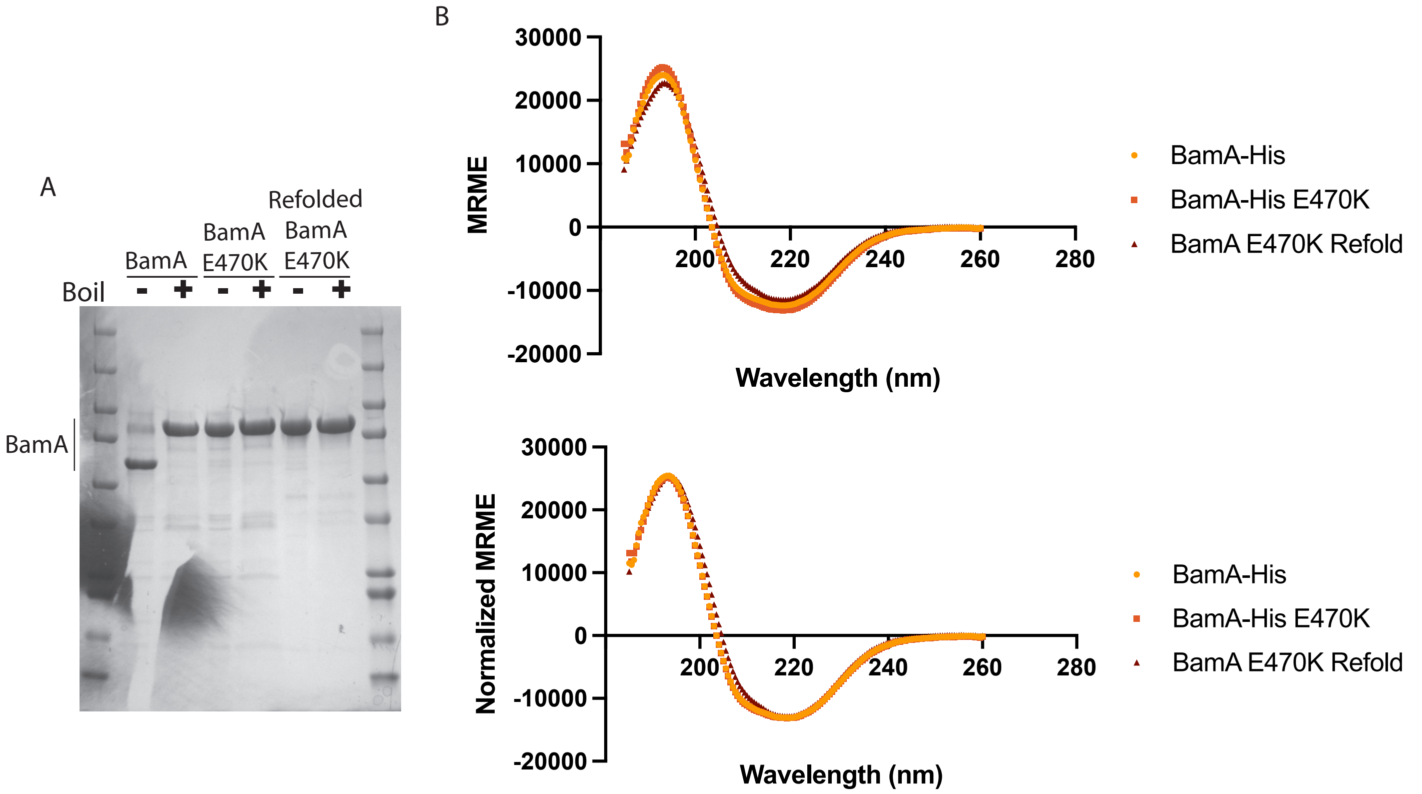


Fig. S6. Refolded BamA E470K adopts a similar structure to membrane-extracted BamA.
(A) SDS-PAGE of purified BamA-His, BamA-His E470K, and refolded BamA E470K. (B) Circular dichroism spectra of protein samples from (A) reported as the mean residue molar ellipticity (MRME) or the Normalized MRME. Curves were normalized to BamA-His E470K by multiplication by a factor of 1.06 and 1.12 for BamA-His and refolded BamA E470K, respectively.

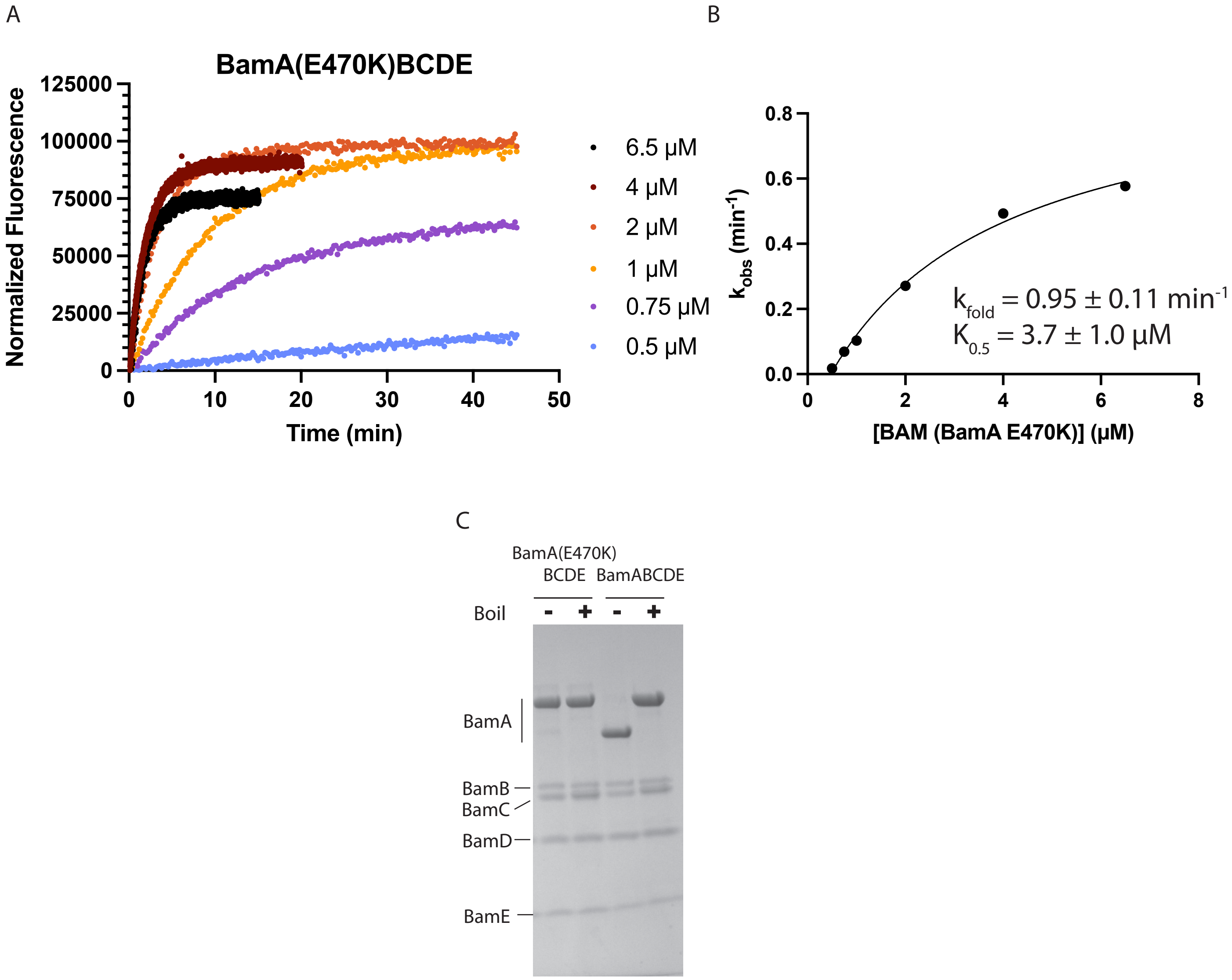


Fig. S7. Activity assay of BAM (BamA E470K) shows activity comparable to WT BAM.
(A) Fluorescence data from one independent single-turnover activity assay of the BAM (BamA E470K) mutant at different concentrations. (B) Single exponential rate constants from (A) were plotted against BAM concentration and fitted to Eq. **Error! Reference source not found.** and represents one biological replicate. Inset values for $k_{fold}$ and $K_{0.5}$ represent the e stimated value ± SE of the fit. (C) SDS-PAGE of purified BamA(E470K)BCDE compared to wildtype BamABCDE.

Table S1
Plasmids used in this study.

| Plasmid | Internal Code | Description | Reference |
|---|---|---|---|
| tOmpA | pMS 471 | pET28a (+) vector containing residues 22-197 of *E. coli* OmpA | This study |
| tOmpA 1 Cys N-term | pMS 755 | tOmpA plasmid with a cysteine introduced at the third position | This study |
| SurA | pMS 332 | pET vector containing WT *E. coli* SurA gene | This study |
| WT BAM | pMS 1289 | pJH114, plasmid with genes encoding all five subunits of the *E. coli* BAM complex | (1) |
| BAM (BamA ∆P1) | pMS 1843 | WT BAM plasmid with BamA gene missing codons V26-R91 | This study |
| BAM (BamA ∆P1-2) | pMS 1844 | WT BAM plasmid with BamA gene missing codons V26-E171 | This study |
| BAM (BamA ∆P1-3) | pMS 1845 | WT BAM plasmid with BamA gene missing codons V26-I260 | This study |
| BamA-His | pMS 1846 | WT BAM plasmid with an N-terminally 6-His tagged BamA gene (coding sequence identical to BamA in complementation plasmid (2)), and deletion of BamBCDE lipoproteins | This study |
| BamA | pMS 1224 | Plasmid with gene encoding mature *E. coli* BamA (codons E22-W810) for expression into inclusion bodies | (3) |
| BamA E470K | pMS 1840 | BamA plasmid (pMS 1224) with BamA E470K point mutation | This study |
| BAM (BamA E470K) | pMS 1841 | WT BAM plasmid with BamA gene with E470K point mutation | This study |
| BamA-His E470K | pMS 1847 | WT BAM plasmid with an N-terminally 6-His tagged BamA gene with E470K point mutation, and deletion of BamBCDE lipoproteins | This study |

Table S2
Oligonucleotides used in this study.

| Oligonucleotide | Sequence (5' → 3') | Description |
|---|---|---|
| U703 | GATATACCATGGCTTGCCCGA | Upper primer for site-directed mutagenesis to introduce an N-terminal cysteine into the tOmpA gene |
| L703 | GGTGTTATCTTTCGGGCAAGC | Lower primer for site-directed mutagenesis to introduce an N-terminal cysteine into the tOmpA gene |
| BamA ∆POTRA1 gene fragment | cgtctcactggtgaaaagaaaaaccaccctggcgcccaatacgcaaaccgcctctccccgcgcgttggccgattcattaa tgcagctggcacgacaggtttcccgactggaaagcgggcagtgagcgcaacgcaattaatgtgagttagcgcgaattga tctggtttgacagcttatcatcgactgcacggtgcaccaatgcttctggcgtcaggcagccatcggaagctgtggtatggctg tgcaggtcgtaaatcactgcataattcgtgtcgctcaaggcgcactcccgttctggataatgttttttgcgccgacatcataac ggttctggcaaatattctgaaatgagctgttgacaattaatcatccggctcgtataatgtgtggaattgtgagcggataacaat ttcacacaggaaacagcaTATGGTTAGGAAGAACGCATAATAACGATGGCGATGAAAAAGTT GCTCATAGCGTCGCTGCTGTTTAGCAGCGCCACCGTATACGGTGCTGAAGGGTTCG TACCGACCATTGCCAGCATTACTTTCTCCGGTAACAAATCGGTGAAAGATGACATGC TGAAGCAAAACCTCGAGGCTTCTGGTGTGCGTGTGGGCGAATCCCTCGATCGCAC CACCATTGCCGATATCGAGAAAGGTCTGGAAGACTTCTACTACAGCGTCGGTAAATA TAGCGCCAGCGTAAAAGCTGTCGTGACCCCGCTGCCGCGCAACCGTGTTGACCTA AAACTGGTGTTCCAGGAAGGTGTGTCAGCTGAAATCCAGCAAATTAACATTGTTGG TAACCATGCTTTCACCACCGACGAACTGATCTCTCATTTCCAACTGCGTGACGAAGT GCCGTGGTGGAACGTGGTAGGCGATCGTAAATACCAGAAACAGAAACTGGCGGGC GACCTTGAAACCCTGCGCAGCTACTATCTGGATCGCGGTTATGCCCGTTTCAACAT CGACTCTACCCAGGTCAGTCTGACGCCAGATAAAAAAGGTATTTACGTCACGGTGA ACATCACCGAAGGCGATCAGTACAAGCTTTCTGGCGTTGAAGTGAGCGGCAACCTT | Gene fragment used in Gibson assembly reaction to substitute wildtype BamA gene with a BamA gene missing POTRA1 domain |

| | | |
|---|---|---|
| | GCCGGGCACTCCGCTGAAATTGAGCAGCTGACTAAGATCGAGCCGGGTGAGCTGTATAACGGCACCAAAGTGACCAAGATGGAAGATGACATCAAAAAGCTT | |
| BamA ΔPOTRA1-2 gene fragment | cgtctcactggtgaaaagaaaaaccaccctggcgcccaatacgcaaaccgcctctccccgcgcgttggccgattcattaatgcagctggcacgacaggtttcccgactggaaagcgggcagtgagcgcaacgcaattaatgtgagttagcgcgaattgatctggtttgacagcttatcatcgactgcacggtgcaccaatgcttctggcgtcaggcagccatcggaagctgtggtatggctgtgcaggtcgtaaatcactgcataattcgtgtcgctcaaggcgcactcccgttctggataatgttttttgcgccgacatcataacggttctggcaaatattctgaaatgagctgttgacaattaatcatccggctcgtataatgtgtggaattgtgagcggataacaatttcacacaggaaacagcaTATGGTTAGGAAGAACGCATAATAACGATGGCGATGAAAAAGTTGCTCATAGCGTCGCTGCTGTTTAGCAGCGCCACCGTATACGGTGCTGAAGGGTTCGTAGGTGTGTCAGCTGAAATCCAGCAAATTAACATTGTTGGTAACCATGCTTTCACCACCGACGAACTGATCTCTCATTTCCAACTGCGTGACGAAGTGCCGTGGTGGAACGTGGTAGGCGATCGTAAATACCAGAAACAGAAACTGGCGGGCGACCTTGAAACCCTGCGCAGCTACTATCTGGATCGCGGTTATGCCCGTTTCAACATCGACTCTACCCAGGTCAGTCTGACGCCAGATAAAAAAGGTATTTACGTCACGGTGAACATCACCGAAGGCGATCAGTACAAGCTTTCTGGCGTTGAAGTGAGCGGCAACCTTGCCGGGCACTCCGCTGAAATTGAGCAGCTGACTAAGATCGAGCCGGGTGAGCTGTATAACGGCACCAAAGTGACCAAGATGGAAGATGACATCAAAAAGCTT | Gene fragment used in Gibson assembly reaction to substitute wildtype BamA gene with a BamA gene missing POTRA1-2 domains |
| BamA ΔPOTRA1-3 gene fragment | cgtctcactggtgaaaagaaaaaccaccctggcgcccaatacgcaaaccgcctctccccgcgcgttggccgattcattaatgcagctggcacgacaggtttcccgactggaaagcgggcagtgagcgcaacgcaattaatgtgagttagcgcgaattgatctggtttgacagcttatcatcgactgcacggtgcaccaatgcttctggcgtcaggcagccatcggaagctgtggtatggctgtgcaggtcgtaaatcactgcataattcgtgtcgctcaaggcgcactcccgttctggataatgttttttgcgccgacatcataacggttctggcaaatattctgaaatgagctgttgacaattaatcatccggctcgtataatgtgtggaattgtgagcggataacaatttcacacaggaaacagcaTATGGTTAGGAAGAACGCATAATAACGATGGCGATGAAAAAGTTGCTCATAGCGTCGCTGCTGTTTAGCAGCGCCACCGTATACGGTGCTGAAGGGTTCGTAACCGAAGGCGATCAGTACAAGCTTTCTGGCGTTGAAGTGAGCGGCAACCTTGCCGGGCACTCCGCTGAAATTGAGCAGCTGACTAAGATCGAGCCGGGTGAGCTGTATAACGGCACCAAAGTGACCAAGATGGAAGATGACATCAAAAAGCTT | Gene fragment used in Gibson assembly reaction to substitute wildtype BamA gene with a BamA gene missing POTRA1-3 domains |
| BamA-His gene fragment | CAATCCGCCCTCACTACAACCGcgtctcactggtgaaaagaaaaaccaccctggcgcccaatacgcaaaccgcctctccccgcgcgttggccgattcattaatgcagctggcacgacaggtttcccgactggaaagcgggcagtgagcgcaacgcaattaatgtgagttagcgcgaattgatctggtttgacagcttatcatcgactgcacggtgcaccaatgcttctggcgtcaggcagccatcggaagctgtggtatggctgtgcaggtcgtaaatcactgcataattcgtgtcgctcaaggcgcactcccgttctggataatgttttttgcgccgacatcataacggttctggcaaatattctgaaatgagctgttgacaattaatcatccggctcgtataatgtgtggaattgtgagcggataacaatttcacacaggaaacagcaTATGGTTAGGAAGAACGCATAATAACGATGGCGATGAAAAAGTTGCTCATAGCGTCGCTGCTGTTTAGCAGCGCCACCGTATACGGTGCTCACCATCACCATCACCATGCTGCAGAAGGGTTCGTAGTGAAAGATATTCATTTCGAAGGCCTTCAGCGTGTCGCCGTTGGTGCGGCCCTCCTCAGTATGCCGGTGCGCACAGGCGACACGGTTAATGATGAAGATATCAGTAATACCATTCGCGCTCTGTTTGCTACCGGCAACTTTGAGGATGTTCGCGTCCTTCGTGATGGTGATACCCTTCTGGTTCAGGTAAAAGAACGTCCGACCATTGCCAGCATTACTTTCTCCGGTAACAAATCGGTGAAAGATGACATGCTGAAGCAAAACCTCGAGGCTTCTGGTGTGCGTGTGGGCGAATCCCTCGATCGCACCACCATTGCCGATATCGAGAAAGGTCTGGAAGACTTCTACTACAGCGTCGGTAAATATAGCGCCAGCGTAAAAGCTGTCGTGACCCCGCTGCCGCGCAACCGTGTTGACCTAAAACTGGTGTTCCAGGAAGGTGTGTCAGCTGAAATCCAGCAAATTAACATTGTTGGTAACCATGCTTTCACCACCGACGAACTGATCTCTCATTTCCAACTGCGTGACGAAGTGCCGTGGTGGAACGTGGTAGGCGATCGTAAATACCAGAAACAGAAACTGGCGGGCGACCTTGAAACCCTGCGCAGCTACTATCTGGATCGCGGTTATGCCCGTTTCAACATCGACTCTACCCAGGTCAGTCTGACGCCAGATAAAAAAGGTATTTACGTCACGGTGAACATCACCGAAGGCGATCAGTACAAGCTTTCTGGCGTTGAAGTGAGCGGCAACCTTGCCGGGCACTCCGCTGAAATTGAGCAGCTGACTAAGATCGAGCCGGGTGAGCTGTATAACGGCACCAAAGTGACCAAGATGGAAGATGACATCAAAAAGCTTCTCGGTCGCTATGGTTATGCCTATCCGCGCGTACAGTCGATGCCCGAAATTAACGATGCCGACAAAACCGTTAAATTACGTGTGAACGTTGATGCGGGTAACCGTTTCTACGTGCGTAAGATCCGTTTTGAAGGTAACGATACCTCGAAAGATGCCGTCCTGCGTCGCGAAATGCGTCAGATGGAAGGTGCATGGCTGGGGAGCGATCTGGTCGATCAGGGTAAGGAGCGTCTGAATCGTCTGGGCTTCTTTGAAACTGTCGATACCGATACCCAACGTGTTCCGGGTAGCCCGGACCAGGTTGATGTCGTCTACAAGGTAAAAGAGCGCAACACCGGTAGCTTCAACTTTGGTATTGGTTACGGTACTGAAAGTGGCGTGAGCTTCCAGGCTGGTGTGCAGCAGGATAACTGGTTAGGTACAGGTTATGCTGTTGGTATCAACGGGACCAAAAACGATTACCAGACCTATGCTGAACTGTCGGTAACCAACCCGTACTTCACCGTAGATGGCGTAAGCCTCGGTGGTCGTCTCTTCTATAATGACTTCCAGGCAGATGACGCCGACCTGTCCGACTATACCAACAAGAGTTATGGTACAGACGTGA | Gene fragment containing an N-terminal 6-His tagged BamA flanked at the end by an XbaI site to exclude BamBCDE when used with restriction cloning |

| | | |
|---|---|---|
| | CGTTGGGCTTCCCGATTAACGAATATAACTCGCTGCGTGCAGGTCTGGGTTATGTAC ATAACTCCCTGTCCAACATGCAGCCTCAGGTTGCGATGTGGCGTTATCTGTACTCTA TGGGTGAACATCCGAGCACCTCTGATCAGGATAACAGCTTCAAAACGGACGACTTC ACGTTCAACTATGGTTGGACCTATAACAAGCTTGACCGTGGTTACTTCCCGACAGAT GGTTCACGTGTCAACCTGACCGGTAAAGTGACCATTCCTGGATCGGATAACGAATA CTACAAAGTGACGTTAGACACGGCGACTTATGTGCCGATCGATGACGATCACAAAT GGGTTGTTCTGGGGCGTACCCGCTGGGGTTATGGTGATGGTTTAGGCGGCAAAGA GATGCCGTTCTACGAGAACTTCTATGCCGGTGGTTCCAGCACCGTGCGTGGCTTCC AGTCCAATACCATTGGTCCGAAAGCAGTTTACTTCCCGCATCAGGCCAGTAATTATG ATCCGGACTATGATTACGAATGTGCGACTCAGGACGGCGCGAAAGACCTGTGTAAA TCGGATGATGCTGTAGGCGGTAACGCCATGGCGGTTGCCAGCCTCGAGTTCATCA CCCCGACGCCGTTTATTAGCGATAAGTATGCTAACTCGGTTCGTACTTCCTTCTTCT GGGATATGGGTACCGTTTGGGATACAAACTGGGATTCCAGCCAATATTCTGGATATC CGGACTATAGTGATCCAAGCAATATCCGTATGTCTGCGGGTATCGCATTACAATGGAT GTCCCCATTGGGGCCGTTGGTGTTCTCCTACGCCCAGCCGTTCAAAAAGTACGATG GAGACAAGGCAGAACAGTTCCAGTTTAACATCGGTAAAACCTGGTAAGTGGGATCT GAGAGGGACCCGtctagaATGCAATTGCGTAAATTACT | |
| U2804 | TATGCTaaaCTGTCGGTAACCAACCCGTACTTCACCGT | Upper primer for site-directed mutagenesis for production of pMS 1840 |
| L2804 | CGACAGtttAGCATAGGTCTGGTAATCGTTTTTGGTCCCGTTGA | Lower primer for site-directed mutagenesis for production of pMS 1840 |

### SI Materials and Methods

*Cloning of tOmpA 1 Cys N-term*
An N-terminal Cys was introduced into tOmpA (residues 22-197 of WT full length OmpA). Plasmids and primers are provided in Tables S1 and S2. This resulted in a construct of mature tOmpA 1 Cys N-term without a signal sequence for expression into inclusion bodies

*Expression of Unfolded OMP Inclusion Bodies*
For expression into inclusion bodies, BL21 (DE3) cells were transformed with plasmid containing tOmpA 1 Cys N-term gene and plated onto LB agar plates supplemented with 50 µg/mL kanamycin. Colonies were used to inoculate 5 mL of LB supplemented with kanamycin and grown overnight at 37°C. 1.25 mL of the overnight starter culture was used to inoculate 100 mL of fresh LB supplemented with kanamycin and cells were grown at 37°C until the culture reached an $OD_{600}$ of 0.4-0.6. Protein expression was induced with 1 mM isopropyl β-D-thiogalactopyranoside (IPTG) and incubated at 37°C for another 3 hours. Cells were harvested by centrifugation and pellets were resuspended in 30 mL 50 mM Tris pH 8, 0.5 mg/mL lysozyme. Cell solution was frozen at -70°C until use.

*Purification of Unfolded OMP from Inclusion Bodies*
To purify unfolded OMP, frozen cell solutions were thawed in warm water bath and benzonase nuclease was added to reduce viscosity. Solutions were sonicated at 70% power for pulses of 5 seconds with 30 second breaks for a total pulse time of 2 minutes. Insoluble material was pelleted by centrifugation at 7650g for 30 minutes at 4°C and pellets were resuspended in 30 mL 50 mM Tris pH 8, 0.5% Triton X-100. Resuspensions were sonicated at 40% power for pulses of 10 seconds with 20 second breaks for a total pulse time of 2 minutes. Insoluble material was pelleted by another round of centrifugation. Pellets were resuspended in 30 mL 10 mM Tris pH 8, 1 mM ethylenediaminetetraacetic acid (EDTA) and pelleting and resuspension was repeated 3 times to remove residual detergent. On the last resuspension, pellet was resuspended in 1.25 mL EDTA buffer supplemented with 1 mM phenylmethylsulfonyl fluoride (PMSF), insoluble material was pelleted again, supernatant was discarded, and pellets were placed at -20°C until use. To prepare solubilized denatured OMP, the purified inclusion body pellet was resuspended in 1.5 mL 6 M guanidine-HCl, 25 mM Tris pH 8, 2 mM TCEP, and solution was filtered with a 0.1 µm filter. Protein solution was snap-frozen in liquid nitrogen and stored at -70°C until use.

*Preparation of Lipid Films*
Phosphatidylcholine (PC) 10:0 (850325, Avanti Research), PC 12:0 (850335, Avanti Research), PC 18:1 (Δ9-cis) (DOPC, 850375, Avanti Research), and *E. coli* Extract Polar (100600, Avanti Research) lipids were purchased as 25 mg/mL stocks of lipids in chloroform. For pure lipid films, aliquoted appropriate volume of lipid stock into glass vial to obtain 10 mM or 8 mg/mL lipids for PC or *E. coli* lipids, respectively, when resuspended in a volume of 250 µL. Lipids were dried under a flow of nitrogen gas to form a thin lipid film, and residual chloroform was evaporated off by vacuum desiccation overnight. Dried lipid films were stored at -20°C under nitrogen gas. For PC 10:0/PC 12:0 mixtures, appropriate volumes of pure chloroform stock were added to form 3:1, 1:1, or 1:3 molar ratios of PC 10:0 to PC 12:0 such that the total lipid concentration when resuspended in 250 µL is 10 mM. Films were formed and dried as before.

*Preparation of PC Liposomes*
To make PC liposomes, rehydrate lipid films in 250 µL 1X Tris-Buffered Saline (TBS) + EDTA (25 mM Tris pH 8, 150 mM NaCl, 1 mM EDTA) at 21°C for 30 minutes to make 10 mM lipid solutions. Solutions were bath sonicated for 15 minutes, allowed to rest for 10 minutes, vortexed at power 6 for 10 seconds, then extruded through a 0.2 µm filter at least 21 times using a mini-extruder (Avanti Research).

*Fluorescence Measurements of tOmpA-A488*
All solution fluorescence measurements were performed with the PTI (Photon Technology International) steady-state spectrofluorometer QuantaMaster (Model QM-6) (Horiba) controlled with the PTI FeliX32 Version 1.2 (2001-2005) software. The instrument was set to a PMT of 1000, excitation wavelength of 485 nm, emission wavelength of 516 nm, 3 nm monochromator slit widths, and a circulating water bath set to 21°C. Real time corrections were enabled and source gain was referenced to 485 nm and set to 1 V. Folding assays were prepared, as described above, and placed in 10 mm rectangular quartz cuvettes (3-Q-10, Starna Cells, Inc) with stir bars. Background measurements were taken for all folding reaction solutions before OMP was added. Folding reactions were initiated by addition of tOmpA-A488 and fluorescence measurements were taken at appropriate times. Processing of the data was performed as described in SI Materials and Methods and data were plotted as the absolute background subtracted fluorescence or the change in fluorescence against time. Curves were fitted to double or single exponential functions with Prism 10 software.

*Quantification of Folding of tOmpA-A488 by SDS-PAGE*
To analyze extent of folding of labeled tOmpA, samples from folding reactions described above were quenched with 4X SDS-PAGE loading buffer (200 mM Tris-HCl, 8% SDS, 6 mM bromophenol blue, 4.3 M glycerol, 0.57 mM 2-mercaptoethanol) and kept on ice until ready to run on gel. For time courses, two samples were taken at the last timepoint,

and one was boiled at 95C for 5 min to re-denature the protein. All samples were loaded onto 12% (w/v) Tris-HCl SDS-PAGE gels and ran at 200V on ice. Gels were imaged using an Amersham Typhoon 5 (Cytiva) in fluorescence mode under a 488 nm laser with a Cy2 filter, a pixel size of 50 or 100 μm, and the PMT set to 465 V. Progression of folding was quantified as an increase in the folded tOmpA-A488 band intensity using a modified protocol with the ImageJ software (Fiji). Briefly, equal-sized boxes were drawn around the entire lane for each sample in the gel. Within the Analyze->Gels->Plot Lanes function, a baseline was drawn for the total lane fluorescence, and vertical lines were drawn to delimit the folded band peak. Only the folded band intensity was considered to eliminate complications from previously observed higher molecular weight aggregates, which would contribute to a loss in the unfolded band intensity (4). Folding was quantified by normalizing the fluorescence of the folded band peak to the total lane fluorescence and data were plotted as the change in normalized folded band intensity from the first time point against time. Folding data were fitted to a single exponential function with Prism 10 software.

*Post-Processing of tOmpA-A488 Fluorescence Measurements*
For PC liposome reactions: all time points were adjusted for the delay in reading the first timepoint after the reaction was initiated, 6 points for each time point (collected as intervals of 2 points/second over 3 seconds) from measurements were averaged (except for the continuously read, PC 10:0 condition), and background fluorescence values were subtracted from each measurement. The initial timepoint fluorescence was subtracted from each timepoint to obtain a change in fluorescence.
For BAM proteoliposome reactions: all time points were adjusted for the delay in reading the first timepoint after the reaction was initiated, and background fluorescence values were subtracted from each measurement. Reaction curves were fitted to single exponential functions in Prism 10 and outliers were detected and eliminated from the fit using the default parameters (Q = 1%). The predicted y-intercept was used to normalize the values to get an estimate of the fluorescence change from time = 0.

*Expression and Purification of SurA*
For expression of SurA, *E. coli* Rosetta (DE3) cells were transformed with a plasmid coding for the SurA protein with an N-terminal 6-histidine tag (pMS 332) and plated onto LB agar supplemented with 50 μg/mL kanamycin. A colony was used to inoculate a 50 mL LB + kanamycin (50 μg/mL) culture and cells were grown overnight at 37°C with shaking. 1L cultures of LB + kanamycin (50 μg/mL) were inoculated with the overnight culture and allowed to grow at 37°C until reaching an $OD_{600}$ of 0.6. 1 mM IPTG was added to cultures to induce expression, and cells were incubated for 5 hours at 37°C. Cells were harvested by centrifugation at 4,000g for 20 minutes. Cell pellets were resuspended in lysis buffer (25 mM Tris pH 7.5, 300 mM NaCl) supplemented with a protease inhibitor cocktail (EDTA acid-free, Roche) and stored at -80°C overnight. Cells were thawed and lysed by sonication with 10 second pulses for a total pulse time of 2 minutes. Lysis was spun at 16,000g for 30 minutes to pellet cell debris. The supernatant was loaded onto a Ni-NTA column equilibrated with lysis buffer. The column was washed with one column volume of lysis buffer, then 10 column volumes of wash buffer (25 mM Tris pH 7.5, 300 mM NaCl, 20 mM imidazole). The protein was eluted from the column with elution buffer (25 mM Tris pH 7.5, 300 mM NaCl, 250 mM imidazole), and fractions containing the protein were pooled.

*Cloning of BAM Complex Mutants*
The parent plasmid containing all the BAM complex genes (BamABCDE-8His) pJH114 was kindly provided by Harris Bernstein and used for subsequent cloning. Plasmids and primers/gene fragments used are provided in Tables S1 and S2. For POTRA deletion mutants, gene fragments were ordered from Twist Bioscience with sequences overlapping BamA, excluding codons for V26 through R91 (for ΔP1), V26 through E171 (for ΔP1-2), or V26 through I260 (for ΔP1-3). Gibson assembly was used to combine these gene fragments with the WT BAM plasmid backbone (created by digestion with restriction enzymes KasI and AleI) For construction of BamA-His plasmid, a gene fragment was ordered from Twist Bioscience covering all the BamA gene of the WT BAM plasmid, with a 6xHis2xAla tag inserted at amino acid position 22 and a XbaI restriction site after the BamA gene. This fragment was digested with restriction enzymes KasI and XbaI and ligated into the similarly digested WT BAM plasmid backbone. For the BAM (BamA E470K) mutant, site-directed mutagenesis was performed using primers upper 2804 and lower 2804 against a pET vector containing WT BamA (pMS 1224). The resulting clone sequence was confirmed by sequencing before digesting with restriction enzymes BstBI and NcoI, then ligating into the similarly digested WT BAM plasmid backbone. For the BamA-His E470K mutant, the plasmids for the BamA-His and the BAM (BamA E470K) mutants were digested with restriction enzymes BstBI and NcoI, and the digested BamA E470K insert was ligated into the BamA-His digested backbone. All ligation and Gibson cloning products were cloned into XL-10 Gold competent cells and clones were confirmed by whole plasmid sequencing (Plasmidsaurus Inc).

*Expression and Purification of BAM Complex Mutants*
For expression of BAM complex mutants, BL21 (DE3) cells were transformed with the appropriate plasmid and plated onto LB agar plates supplemented with 100 μg/mL ampicillin. Colonies were used to inoculate 6 mL of LB supplemented with ampicillin and incubated for 7 hours at 37°C. Cells were pelleted, media decanted, and cells were resuspended in fresh LB media. 1 mL of washed cells were used to inoculate 1L of LB supplemented with 100 μg/mL ampicillin and lactose autoinduction supplement (0.6% glycerol, 0.05% glucose monohydrate, 0.2% lactose monohydrate) and cells were grown

overnight with shaking at 37°C. Cells were harvested by centrifugation, pellets were resuspended in 10 mL lysis buffer (50 mM Tris pH 8, 2 mM EDTA, 0.5 mg/mL lysozyme, protease inhibitor cocktail) per 1L culture, and cell solutions were frozen at -70°C. Cell solutions were thawed in a warm water bath and supplemented with Benzonase nuclease, then lysed with an Emulsiflex C3 Homogenizer (Avestin). Solutions were centrifuged at 7650g for 30 min at 4°C to pellet cell debris, then the supernatant was spun at 256,630g for 2 hours at 4°C. The resulting membrane pellet was resuspended in 10 mL extraction buffer (25 mM Tris pH 8, 150 mM NaCl, 1% n-dodecyl-β-D-maltoside (DDM)) per gram of wet membrane and incubated with rocking overnight at 4°C. Extraction was centrifuged again at 256,630g for 1.5 hours at 4°C to pellet insoluble content and the supernatant was filtered to 0.22 μm then affinity purified with a 5 mL HisTrap FF on the AKTA Pure FPLC system. Briefly, the extraction was loaded onto the column equilibrated with Buffer A (25 mM Tris pH 8, 150 mM NaCl, 0.5 mM TCEP, 0.03% DDM) +5% Buffer B (Buffer A + 500 mM imidazole), the column was washed with 4 column volumes 5% Buffer B, then the protein was eluted over a gradient of 5-100% Buffer B over 5 column volumes. Fractions containing protein were pooled, concentrated to ~10 mL (Vivaspin 20 100K MWCO PES concentrator), filtered to 0.22 μm, then gel filtered with a HiLoad 26/600 Superdex 200 pg column in 25 mM Tris pH 8, 150 mM NaCl, 0.5 mM TCEP, 0.03% DDM. Fractions containing protein were pooled and concentrated with a Vivaspin concentrator. Protein concentrations were measured using a NanoDrop with $A_{280}$ measurements. Extinction coefficients were calculated with ProtParam (Expasy) as 291650 $M^{-1}cm^{-1}$ for WT BAM, BAM (BamA ΔP1), and BAM (BamA E470K), 287180 $M^{-1}cm^{-1}$ for BAM (BamA ΔP1-2), 249250 $M^{-1}cm^{-1}$ for BAM (ΔBamC), 205820 $M^{-1}cm^{-1}$ for BAM (BamA ΔP1-3), and 140040 $M^{-1}cm^{-1}$ for BamA-His, BamA-His E470K, and refolded BamA E470K. Protein quality was checked by running purified samples on 4-20% Mini-PROTEAN TGX Protein Gels (Bio-Rad) and staining with Coomassie. Protein stocks were then used immediately or snap frozen with liquid nitrogen and stored at -70°C until use.

*Expression, Purification, and Refolding of Unfolded BamA E470K from Inclusion Bodies*
BamA E470K was expressed into inclusion bodies and purified as described above. To refold BamA E470K, the inclusion body pellet was dissolved in 8M urea, 20 mM Tris pH 8, 1 mM TCEP, filtered with a 0.1 μm filter, and 2 mL of the BamA E470K stock in urea was added slowly to 18 mL 25 mM Tris pH 8, 150 mM NaCl, 1 mM TCEP, 1% DDM with constant stirring at room temperature for 24 hours. To purify refolded BamA E470K, the solution in was spun at 20,000g in a tabletop centrifuge to pellet aggregates. The supernatant was filtered to 0.1 μm, then subjected to size-exclusion chromatography and processed as described above for BAM complex mutants.

*Circular Dichroism of BamA Samples*
To obtain CD spectra of the BamA samples, BamA, BamA E470K, and refolded BamA E470K were diluted into 10 mM phosphate pH 8, 0.03% DDM to final concentrations of 0.05 mg/mL protein in 5 mM NaCl. A blank solution and BamA samples were read on the Applied Photophysics Chirascan Plus CD and Fluorescence Spectrometer in a 1 mm quartz cuvette, measured from 185-200 nm. The pathlength and protein concentration were used to convert measurements to mean residue molar ellipticity (MRME).

*Reconstitution of BAM Complex into E. coli Polar Liposomes*
Briefly, *E. coli* polar lipid (EPL) films were rehydrated with 1X TBS + EDTA buffer (25 mM Tris pH 8, 150 mM NaCl, 1 mM EDTA) supplemented with 0.1% DDM to a concentration of 6 mg/mL EPL, vortexed to assist rehydration, allowed to incubate at room temperature for at least 10 minutes, vortexed again, then extruded through a 0.2 μm filter. EPL solutions were mixed with BAM solutions (maintaining a ratio around 10 μM BAM to 3.33 mg/mL EPL) and dialyzed against 1X TBS + EDTA buffer at room temp for 2 days with a total of 4 buffer exchanges using Slide-A-Lyzer MINI 20K MWCO, 2 mL Dialysis Devices with continuous magnetic stirring in the buffer chamber. Proteoliposomes were extruded first through a 1 μm filter about 5 times, then through a 0.2 μm filter about 7 times to homogenize liposome solution and remove lipid aggregates. This step is crucial to avoiding unwanted fluorescence signal in folding assays due to OMP aggregation at lipid aggregates. Final concentration of BAM proteoliposomes was quantified using a Rapid Gold BCA Protein Assay Kit (Pierce) with BSA as the standard. Molecular weights used to convert to molar concentrations were 199892.73 Da for WT Bam ABCDE, 199891.79 Da for BamA(E470K)BCDE, 192627.5 Da for BamA(ΔP1)BCDE, 183837.49 Da for BamA(ΔP1-2)BCDE, 133775.39 Da for BamA(ΔP1-3)CDE, 89320.05 Da for BamA-His, 89319.11 Da for BamA-His E470K, and 88485.29 Da for folded BamA E470K. Empty EPL liposomes were prepared as above, except instead of adding BAM, 1X TBS + EDTA supplemented with DDM was added to give a final added concentration of DDM of 0.167%, which amounts to 327:1 mole DDM:mole BAM (this is a ballpark ratio for detergent occupancy on membrane proteins (5), and is supported by measurements of the total DDM in a BAM purification, data not included).
